# A diagnostic multiplex PCR scheme for identification of plant-associated bacteria of the genus *Pantoea*

**DOI:** 10.1101/456806

**Authors:** Kossi Kini, Raoul Agnimonhan, Rachelle Dossa, Drissa Silué, Ralf Koebnik

## Abstract

**Background:** The genus *Pantoea* forms a complex of more than 25 species, among which several cause diseases of several crop plants, including rice. Notably, strains of *Pantoea ananatis* and *Pantoea stewartii* have been found to cause bacterial leaf blight of rice in Togo and Benin, while other authors have observed that *Pantoea agglomerans* can also cause bacterial leaf blight of rice. The contribution of these and perhaps other species of *Pantoea* to plant diseases and yield losses of crop plants is currently not well documented, partly due to the lack of efficient diagnostic tools.

**Result:** Using 34 whole genome sequences of the three-major plant-pathogenic *Pantoea* species, a set of PCR primers that specifically detect each of the three species, *P. agglomerans*, *P. ananatis*, and *P. stewartii*, was designed. A multiplex PCR protocol which can distinguish these three species and also detects members of other *Pantoea* species was further developed. Upon validation on a set of reference strains, 609 suspected *Pantoea* strains that were isolated from rice leaves or seeds originating from 11 African countries were screened. In total, 41 *P. agglomerans* strains from eight countries, 79 *P. ananatis* strains from nine countries, 269 *P. stewartii* strains from nine countries and 220 unsolved *Pantoea* strains from ten countries were identified. The PCR protocol allowed detecting *Pantoea* bacteria grown in vitro, in planta and in rice seeds. The detection threshold was estimated at 5 ng/mL of total genomic DNA and 1 × 10^5^ CFU/mL of heated cells.

**Conclusion:** This new molecular diagnostic tool will help accurately diagnose major plant-pathogenic species of *Pantoea*. Due to its robustness, specificity, sensitivity, and cost efficiency it will be very useful for plant protection services and for the epidemiological surveillance of these important crop-threatening bacteria.

## Background

The genus *Pantoea* was first described in 1989 and was recently taxonomically classified as a member of the *Erwiniaceae* family [1]. More than 25 species of this genus have been described and reported worldwide [2,3]. Etymologically, the genus name *Pantoea* is derived from the Greek word ‘Pantoios’, which means “of all sorts or sources” and reflects the diverse geographical and ecological sources from which the bacteria have been isolated. Several species of the genus are qualified as versatile and ubiquitous bacteria because they have been isolated from many different ecological niches and hosts [2,4]. Remarkably, some species have the ability to colonize and interact with members of both the plant and the animal Kingdom [5]. Among the plant-interacting species, *Pantoea ananatis, Pantoea agglomerans* and *Pantoea stewartii* are well known for their phytopathogenic characteristics. They are recognized as the causal agent of several diseases, such as leaf blight, spot disease, dieback, grain discoloration, seed stalk rot, center rot, stem necrosis, palea browning, bulb decay etc. and affect several economically important crops, including cereals, fruits and vegetables [2,6,7].

Bacterial leaf blight caused by *Xanthomonas oryzae* pv. *oryzae* is an important disease of rice and affects rice cultivation in most regions of the world were rice is grown. The bacterium has been associated with this disease since a very long time [8]. Surveys were conducted from 2010 to 2016 to estimate the extent and importance of the disease and the phytosanitary status of rice fields in West Africa. While leaves showing bacterial blight (BB)-like symptoms were frequent, isolation or molecular detection of xanthomonads using the Lang et al diagnostic tool [9] often failed. Instead, other bacteria forming yellow colonies were observed and turned out to belong to the species *P. ananatis* or *P. stewartii*, as documented for samples from Togo and Benin [10,11]. Additionally, other cases of BB and grain discoloration caused by *Sphingomonas* sp. and other undescribed species have been detected in several sub-Saharan Africa countries [12]. This situation represents an “emerging” bacterial species complex that may constitute a threat to rice production in Africa. Therefore, a robust, specific, sensitive, and cost efficient diagnostic tool is of primary importance for accurate pathogen detection. However, none of the several simplex and multiplex PCR tools [13–18] and other molecular [19–24], physiological, biochemical [24–28] diagnostic tools available for *Pantoea* allows accurate simultaneous detection of the three major plant-pathogenic *Pantoea* species. Some of these methods are poorly reproducible and often limited to a single species while others are reproducible but again limited to one species or are not suited to doubtlessly detect African strains.

To overcome this unsatisfying situation, a molecular method was set up for detecting in a single reaction the three major plant-pathogenic *Pantoea* species (*P. ananatis, P. stewartii* and *P. agglomerans*), as well as other members of the genus. A universal multiplex PCR tool was therefore developed and first tested in silico on available genome sequences and on a set of reference strains from USA, Brazil, Spain and Japan. Afterwards, 609 suspected *Pantoea* strains from eleven Africans countries were evaluated with the newly described diagnostic tool. *P. agglomerans* was detected in rice leaves from several African countries for the first time. Finally, the specificity and sensitivity of the multiplex PCR was monitored by analyzing serial dilutions of genomic DNA, serial dilutions of bacterial cell suspensions and solutions of ground leaves and seeds that had been artificially or naturally infected. This new diagnostic tool will prove useful for phytosanitary services in routine diagnostics of *Pantoea* spp in any type of sample (e.g. leaves, seeds, soil, water).

## Materials and Methods

### Bioinformatics prediction of specific PCR primers

*Pantoea* genome sequences were retrieved from NCBI GenBank (Table 1). Sequences for housekeeping genes were identified by TBLASTN [29]. Sequences were then aligned with MUSCLE [30] at EMBL-EBI [31]. Diagnostic primers that can differentiate the three species*, P. agglomerans*, *P. ananatis* and *P. stewartii*, and one primer pair that would amplify DNA from the whole *Pantoea* genus were designed manually. The Tm for PCR primers were automatically predicted by Tm calculator tool at http://www.thermoscientificbio.com/webtools/multipleprimer/ which was developed based on the modified nearest-neighbor interaction method [32].

**Table 1:**
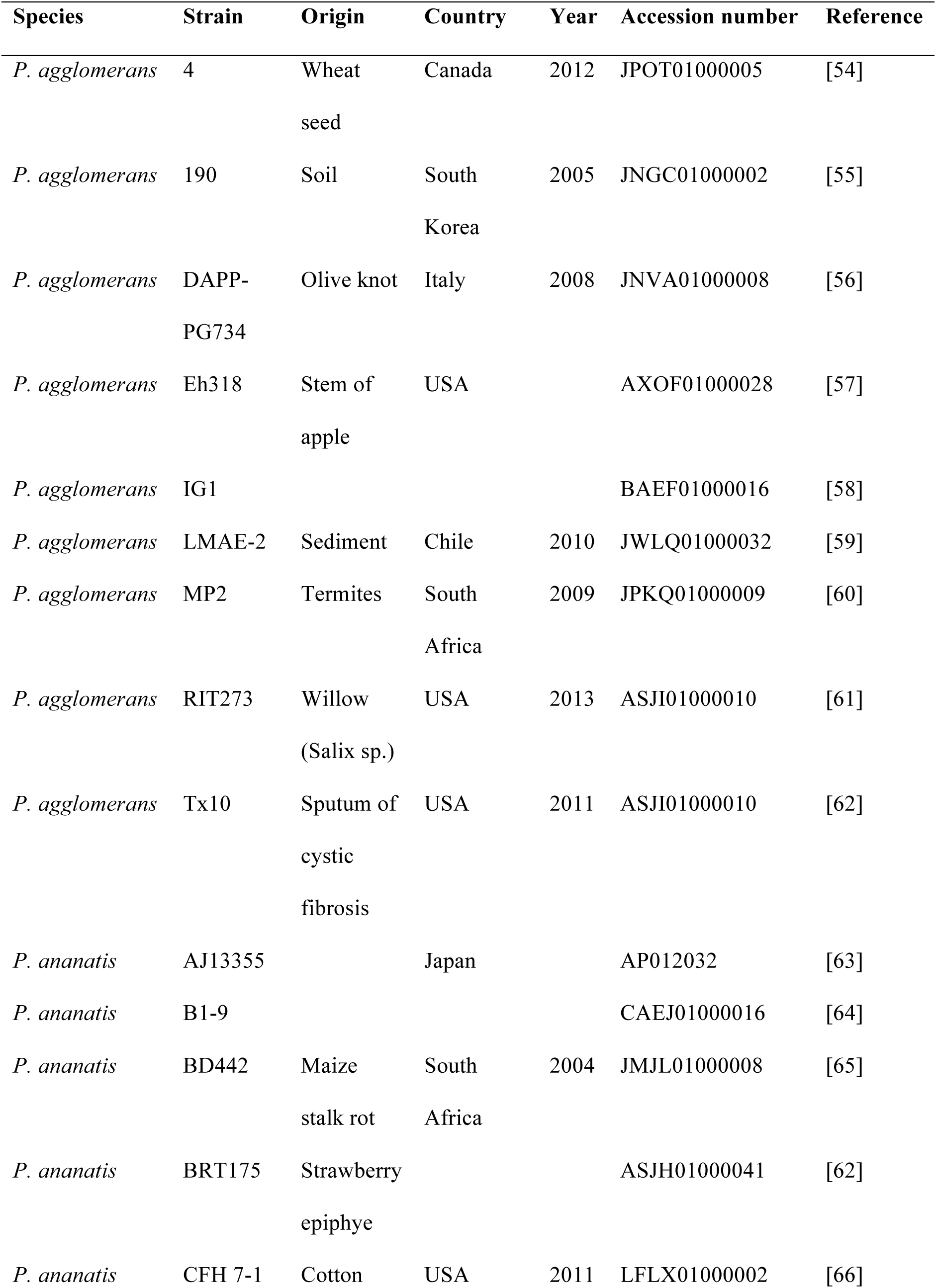
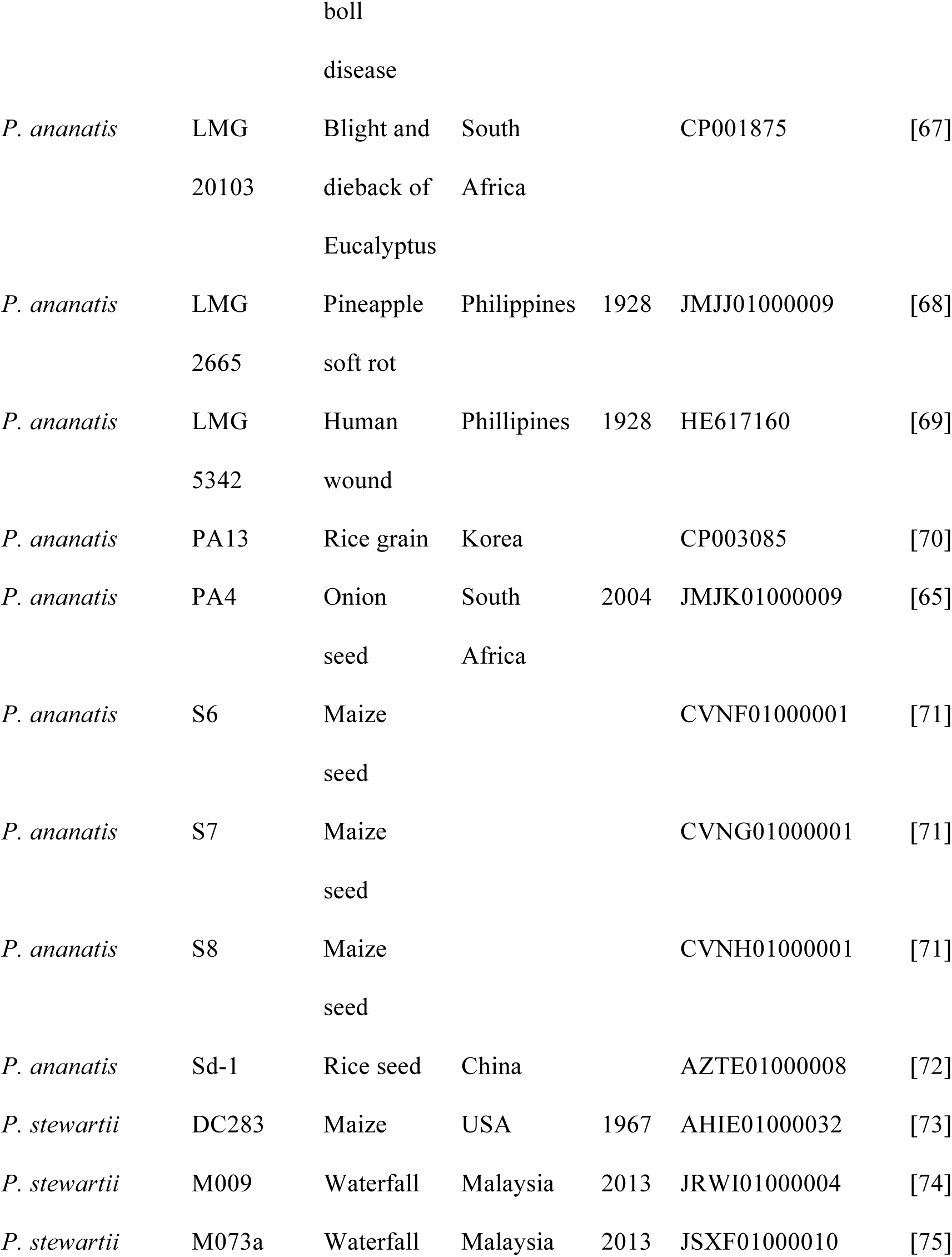
List of *Pantoea* genome sequences used for primers design.

### Optimization of the multiplex PCR

Different types of samples including total genomic DNA, bacterial cells, symptomatic rice leaves, as well as discolored and apparently healthy rice seeds were analyzed. Plant material was ground and macerated before use. To develop a multiplex PCR scheme, individual primer pairs were first tested against the different samples mentioned above, using annealing temperatures close to the predicted Tm (Tm ± 5 °C) and with progressive number of PCR cycles (25 to 35). Primer pairs were then mixed from duplex to quintuplex and PCR conditions were evaluated, testing annealing temperatures close to the optimal Tm of the individual primer pairs (Tm ± 3 °C) and various numbers of PCR cycles. At the end, three promising combinations of annealing temperatures and numbers of PCR cycles were re-evaluated in simplex PCR with the samples mentioned above. The best combination with high specificity and without background amplification was selected as the new diagnostic tool (Tables 2 to 4).

**Table 2:**
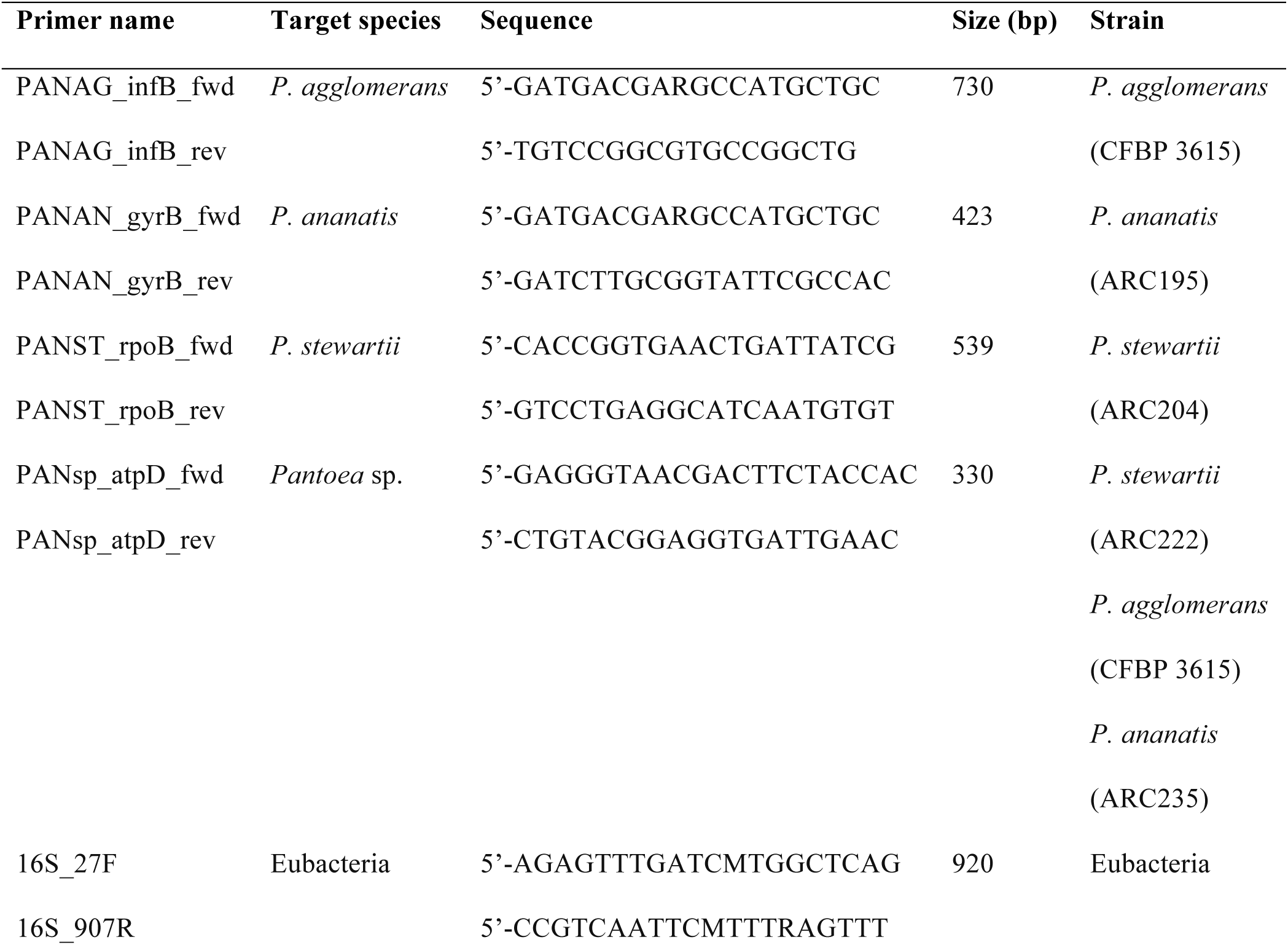
List of PCR primers developed for the *Pantoea* mPCR along with the sequences of the GenBank accessions and the corresponding strains.

**Table 3:** Composition of the multiplex polymerase chain reaction.

| PCR component | Volume per reaction (µL) |  | Final concentration |
| --- | --- | --- | --- |
| Type of template | Purified DNA | Bacterial cells |  |
| Buffer (5x) | 5.0 | 5.0 | 1x |
| dNTPs (2.5 mM each) | 0.5 | 0.5 | 50 µM each |
| Oligonucleotides (10 µM) | 0.4 | 0.4 | 0.16 µM each |
| Takara ExTaq<sup>TM</sup> (5 units/µL) | 0.1 | 0.1 | 0.5 U |
| Template | 2.0 | 5.0 |  |
| Sterile nanopure water<br>(Promega) | 13.4 | 10.4 |  |
| Total | 25.0 | 25.0 |  |

**Table 4:** Reaction parameters of the multiplex PCR thermocycler program.

| Step | Phase | Time | Temperature (°C) |
| --- | --- | --- | --- |
| 1 | Initial denaturation | 3 min | 94 |
| 2 | Denaturation | 30 sec | 94 |
| 3 | Annealing | 30 sec | 58 |
| 4 | Extension | 2 min | 72 |
| 5 | Cycling (steps 2-4) | 30 cycles |  |
| 6 | Final extension | 10 min | 72 |
| 7 | Soak/hold | ∞ | 4-10 |
| 8 | End |  |  |

### Evaluation of the sensitivity of the multiplex PCR scheme using genomics DNA and heat cells

Simplex and multiplex PCR were then used to evaluate the sensitivity of all the species-specific primer pairs individually or in combination with the genus-specific and the 16 sRNA primer pairs. Serial dilutions of total genomic DNA and heated bacterial cells were used for this evaluation. Three *Pantoea* strains, *P. ananatis* strain ARC60, *P. stewartii* strain ARC222, and *P. agglomerans* strain CFBP 3615, were used and distilled sterilized served as a negative control.

To evaluate the PCR scheme on live plant material, leaves and seeds were artificially infected with strains of the three *Pantoea* species. Rice leaves of the cultivar Azucena were inoculated as described previously [10,11]. To produce contaminated seeds, early maturity panicles of the Azucena rice cultivar were spray-inoculated with a 5%-gelatinized bacterial solution (10^6^ CFU/mL). Distillated and gelatinized (5%) sterile water served as a negative control. Three weeks post inoculation, approximately 40% of the grains in the panicles exhibited discolorations. Panicles inoculated with sterile distilled water showed no symptoms. A total of five grains whose surface was first treated with a solution of hypochlorite (10%) and ethanol (70%) and then rinsed with sterile distilled water were ground in 100 mL of sterile distilled water. After centrifugation, the supernatant was used for PCR.

### Evaluation of the multiplex PCR scheme on a large collection of African *Pantoea* strains

Bacterial strains used in this study are listed in Additional file 1. In total, 615 *Pantoea* strains from eleven Africans countries (Benin, Burkina Faso, Burundi, Ghana, Ivory Coast, Mali, Niger, Nigeria, Senegal, Tanzania, Togo) and seven reference strains from USA, Brazil, Spain and Japan were analysed by the new diagnostic tool. The African strains were isolated from rice leaves with BB symptoms, and from discolored and apparently healthy rice seeds. The samples had been collected from 2008 to 2016 in the main rice-growing areas of the countries. Other bacteria, including *Xanthomonas* spp, *Sphingomonas* spp, *Escherichia coli*, *Erwinia* spp, *Burkholderia* spp, and *Pseudomonas* spp, were used as controls. The strains were purified as single colonies, individually grown and preserved as pure cultures following routine methods [33]. Bacterial colonies were grown for 24 to 48 h on PSA plates containing 10 g peptone, 10 g sucrose, 16 g agar and 1 g glutamic acid per liter. Total genomic DNA was extracted using the Wizard genomic DNA purification kit (Promega) according to the manufacturer’s instructions. DNA quality and quantity were evaluated by agarose gel electrophoresis and spectrophotometry (Nanodrop Technologies, Wilmington, DE).

## Results

### Development of a diagnostic PCR scheme for plant-associated *Pantoea*

We aimed at designing diagnostic PCR primers that would target conserved housekeeping genes. The rationale behind was that these genes should be present in all strains, including genetic lineages that have not yet been discovered and would not be present in any strain collection. At the same time, we knew from previous work that sequences of housekeeping genes are divergent enough to doubtlessly distinguish and identify *Pantoea* strains at the species level.

A diagnostic *Pantoea* multiplex PCR method was developed in two steps. First, a complete inventory of publicly available *Pantoea* genome sequences was compiled, consisting of nine *P. agglomerans*, 14 *P. ananatis*, and three *P. stewartii* sequences, totaling to 26 whole genome sequences (Table 1). Complete coding sequences of four housekeeping genes that have previously been used for multilocus sequence analyses (MLSA) of *Pantoea* species [2], *atpD*, *gyrB*, *infB*, and *rpoB*, were then extracted and aligned. Sequence regions that were conserved in all strains of one species but were significantly different in the other two species were identified manually and chosen to design PCR primers (Table 2). To allow multiplexing, we made sure that the amplicon sizes would be between 400 and 750 bp and different enough to be easily distinguishable from each other upon gel electrophoresis (Fig. 1). As a positive control for the PCR reaction, one primer pair was included that would amplify DNA from all bacteria belonging to the *Pantoea* genus, resulting in a smaller amplicon of less than 400 bp. Finally, as a second control, a primer pair was included that targets the ribosomal 16S rRNA gene and leads to an amplicon that is larger than the four *Pantoea*-specific amplicons.

**Figure 1:**
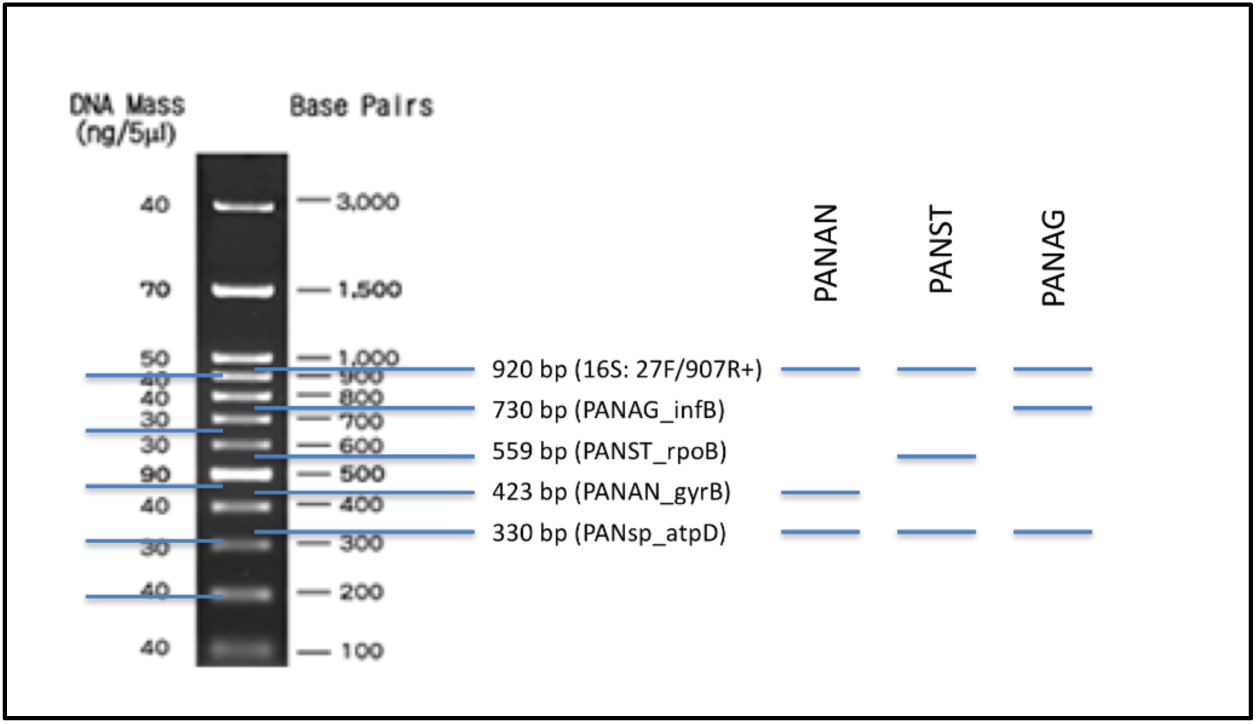
Schematic representation of the multiplex PCR scheme. Sizes of the five expected PCR amplicons are indicated in the middle and their expected migration in a 1.5% TBE agarose gel is shown on the left side. Diagnostic band patterns for the three plant-associated *Pantoea* species are shown on the right side.

In the second step, all primer pairs (Table 2) were evaluated, first by simplex PCR and then by multiplex PCR, with increasing number of primer pairs, as explained in Material and Methods. Three *Pantoea* reference strains were used to develop the PCR scheme using genomic DNA and heat-inactivated bacteria: *P. agglomerans* strain CFBP 3615, *P. ananatis* strain ARC60 and *P. stewartii* strain ARC222 (Fig. 2). Agarose gel electrophoresis demonstrated that the multiplex PCR was able to detect and distinguish all three *Pantoea* species. Notably, the multiplex PCR scheme was also able to detect two or three *Pantoea* species when the corresponding species were present in the same template DNA, as demonstrated by PCR reactions containing equal amounts of DNA of the different species (Fig. 2).

**Figure 2:**
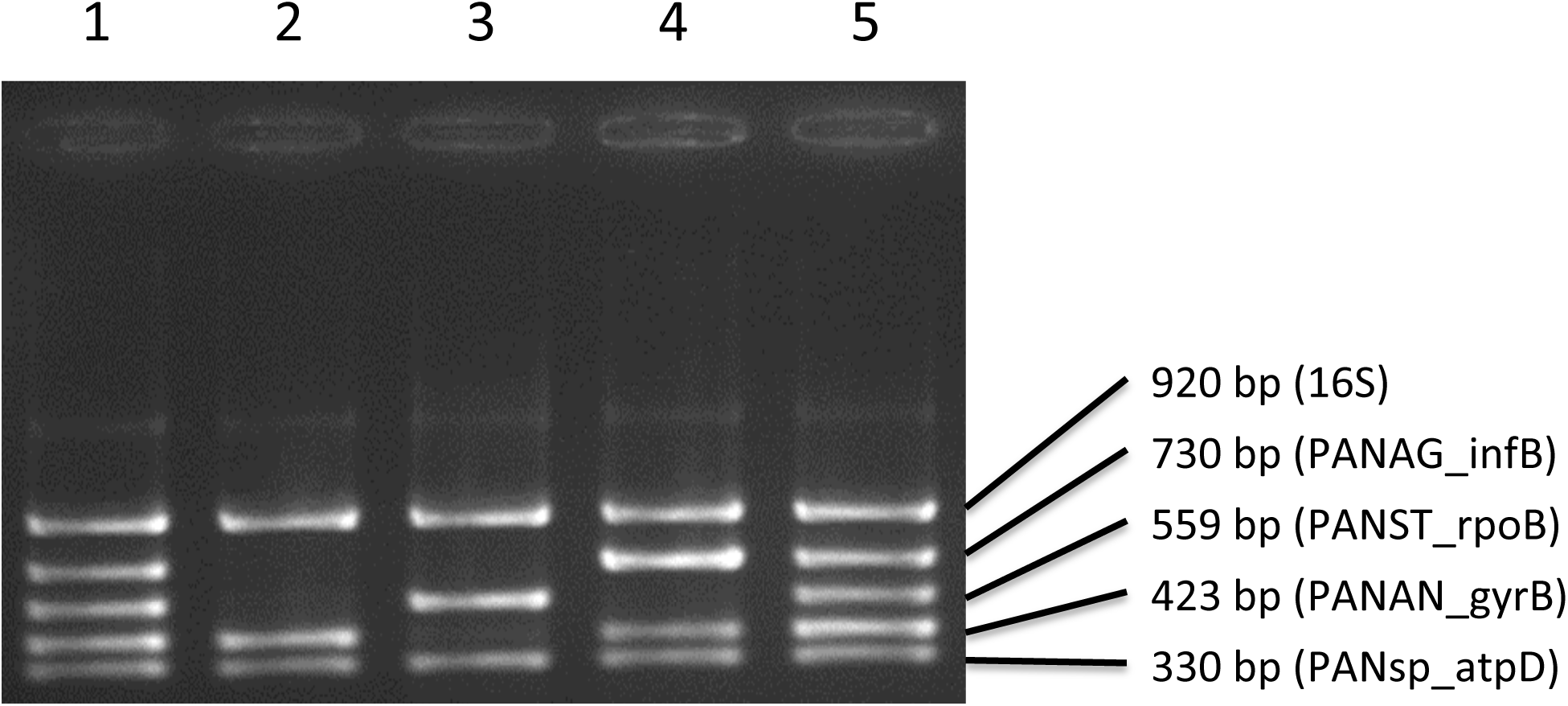
Detection of three *Pantoea* species by multiplex PCR, using heated cell suspensions or genomic DNA as template. Three reference strains were used as representatives for the three *Pantoea* species, *P. ananatis* strain ARC60, *P. stewartii* strain ARC222, and *P. agglomerans* strain CFBP 3615. Lanes 1 & 5, pool of heated cells of the three *Pantoea* species; lane 2, *P. ananatis*; lane 3, *P. stewartii*; lane 4, pool of genomic DNA from *P. ananatis* and *P. agglomerans*;

To simplify the analyses and to avoid isolation of bacteria from plant samples, thus reducing the costs per sample, the PCR scheme was also evaluated on infected leaf material and contaminated seeds. As shown in Fig. 3, the multiplex PCR was able to doubtlessly detect all three *Pantoea* species in both types of plant samples, as demonstrated for the strains CFBP 3615 (*P. agglomerans*), ARC60 (*P. ananatis*), and ARC222 (*P. stewartii*). At the end, a robust PCR protocol was available that was able to amplify DNA from total genomic DNA, bacterial cells, symptomatic rice leaves and from infected rice seeds.

**Figure 3:**
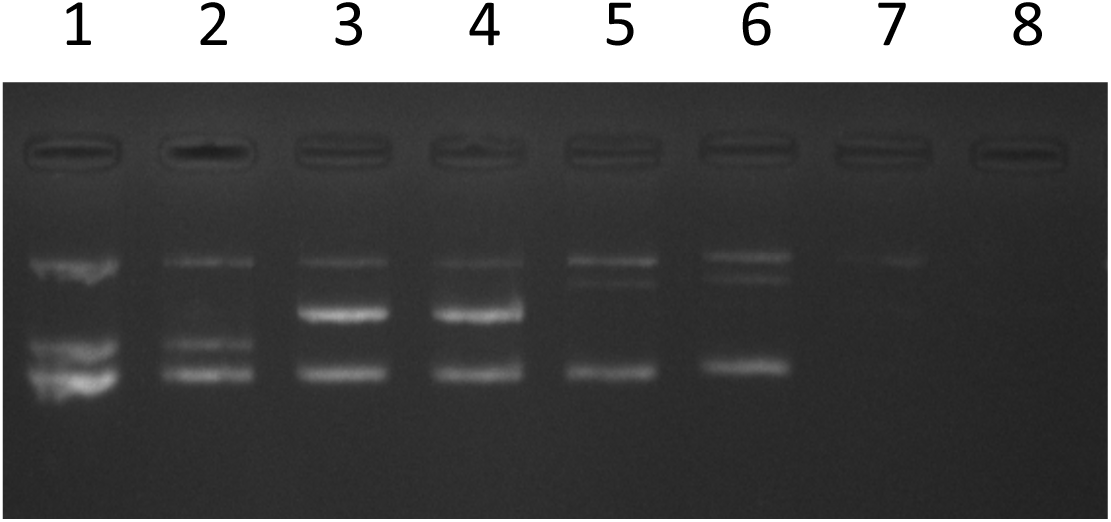
Detection of three *Pantoea* species in artificially infected rice leaves and in contaminated seeds. The following *Pantoea* strains were used: *P. ananatis* strain ARC60, *P. stewartii* strain ARC222, and *P. agglomerans* strain CFBP 3615. Lane 1, *P. ananatis* (leaf sample); lane 2, *P. ananatis* (seed); lane 3, *P. stewartii* (leaf); lane 4, *P. stewartii* (seed); lane 5, *P. agglomerans* (leaf); lane 6, *P. agglomerans* (seed); lane 7, A yellow bacterial colony isolated from rice seeds; lane 8, water.

### Evaluation of the sensitivity of the multiplex PCR scheme using genomic DNA and heated cell suspensions

The evaluation by simplex and multiplex PCR showed that all the species-specific primers were very sensitive individually or in combination with the genus-specific and the 16 sRNA universal primers (Fig. 4). The most sensitive primer pair in simplex PCR was the one targeting *P. stewartii* with a detection limit of 5 pg under our experimental conditions, followed by the *P. agglomerans*-specific primer pair (detection limit of 50 pg) and the *P. ananatis*-specific primer pair (detection limit of 0.5 ng). A similar trend was observed in the multiplex PCR on genomic DNA, with the same detection limit as in simplex PCR for *P. stewartii* and *P. ananatis* and a tenfold less sensitivity for *P. agglomerans*.

**Figure 4:**
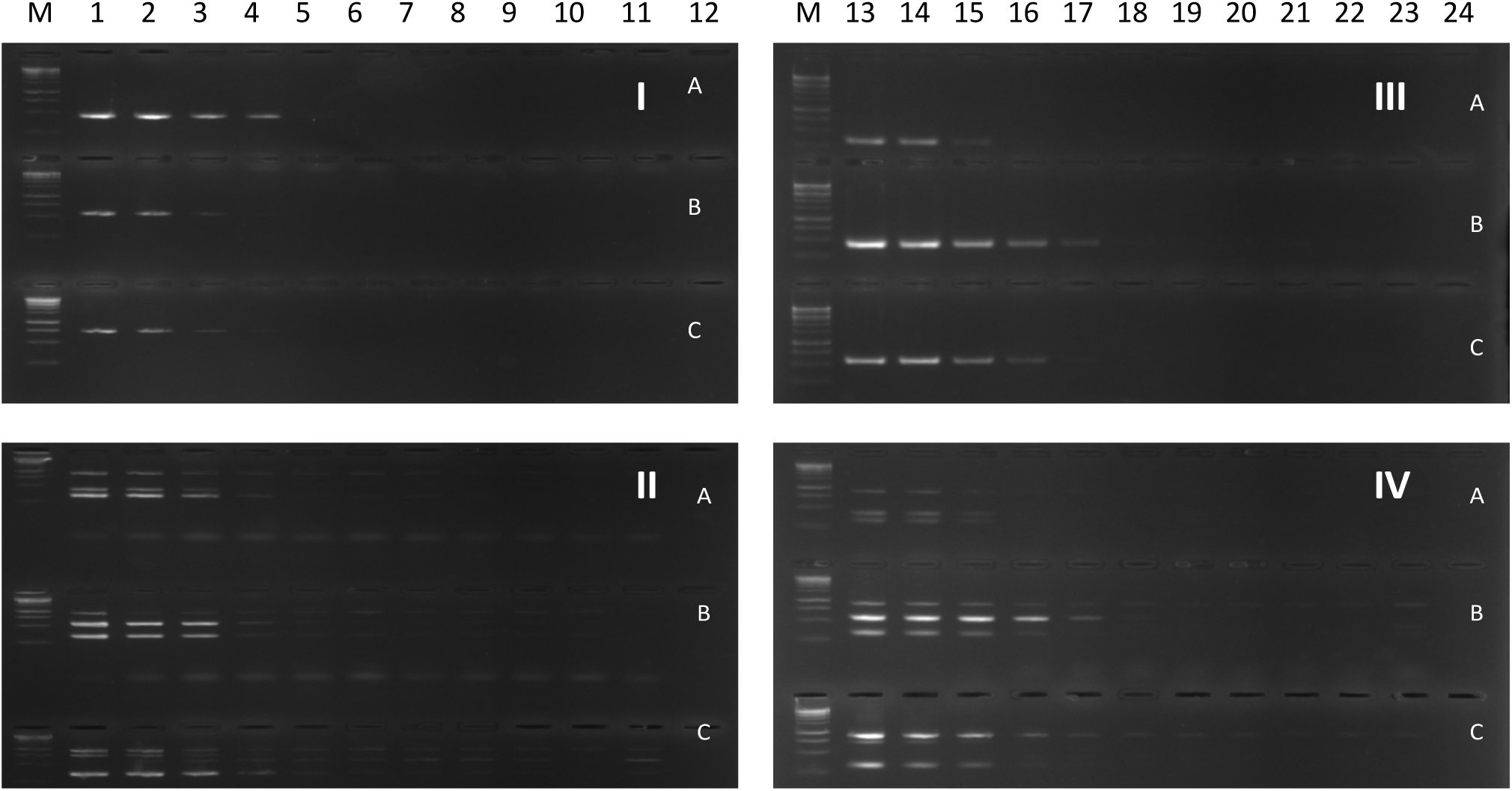
Sensitivity of PCR amplification in simplex and multiplex PCR. Serial dilutions of total genomic DNA and heated bacterial cells were evaluated. I, simplex PCR with bacterial cells; II, multiplex PCR with bacterial cells; III, simplex PCR with genomic DNA; IV multiplex PCR with genomic DNA. Three *Pantoea* strains were used: *P. ananatis* strain ARC60 (A), *P. stewartii* strain ARC222 (B), and *P. agglomerans* strain CFBP 3615 (C). Simplex PCR were performed with the corresponding, species-specific primer pairs, PANAN_gyrB for *P. ananatis*, PANST_rpoB for *P. stewartii*, and PANAG_infB for *P. agglomerans*. The multiplex PCR included all five primer pairs. The following amounts of bacteria or genomic DNA were used as templates for the PCR, corresponding to 10-fold serial dilutions: Lanes 1 to 12 10^6^ CFU/mL, 10^5^ CFU/mL, 10^4^ CFU/mL, 10^3^ CFU/mL, 10^2^ CFU/mL, 10^1^ CFU/mL, 10^0^ CFU/mL, 10^-1^ CFU/mL, 10^-2^ CFU/ml, 10^-3^ CFU/ml, 10^-4^ CFU/mL and water; lanes 12 to 24, 50 ng, 5 ng, 0.5 ng, 50 pg, 5 pg, 0.5 pg, 50 fg, 5 fg, 0.5 fg, 50 ag, 5 ag and water; M, molecular size marker (1 kb DNA ladder, Promega).

When heated bacterial cell suspensions were used as template, the *P. ananatis*-specific primer pair was the most sensitive allowing detection of 10^3^ CFU/mL, while the other two primer pairs were able to detect 10^4^ CFU/mL. However, when all five primers pairs were used in multiplex, the sensitivity was very similar for all three species with a detection limit of approximately 10^4^ CFU/mL.

### Evaluation of the multiplex PCR scheme on a large collection of African *Pantoea* strains

Because recent surveys had indicated that *Pantoea* species could be responsible for many unsolved infections of rice fields in sub-Saharan Africa [10,11], we screened a large collection of isolates. We first re-evaluated a few African strains that had been identified as *P. ananatis* (ARC22, ARC60, ARC651) and *P. stewartii* (ARC229, ARC570, ARC646), using species-specific and the genus-specific PCR primers [10,11]. The multiplex PCR scheme confirmed their previous taxonomic classification. Next, we screened a large collection of African bacterial isolates from rice samples (>1000 strains) among which 609 strains were found to belong to the genus *Pantoea* (Additional file 1). Specifically, this work diagnosed 41 *P. agglomerans* strains from eight countries (Benin, Ghana, Mali, Niger, Nigeria, Senegal, Tanzania, Togo), 79 *P. ananatis* strains from nine countries (Benin, Burkina Faso, Burundi, Mali, Niger, Nigeria, Senegal, Tanzania, Togo), 269 *P. stewartii* strains from nine countries (Benin, Burkina Faso, Ivory Coast, Mali, Niger, Nigeria, Senegal, Tanzania, Togo) and 220 *Pantoea* sp. strains from ten countries (Benin, Burundi, Ghana, Ivory Coast, Mali, Niger, Nigeria, Senegal, Tanzania, Togo) (Additional file 1). This result provided first insights on the presence and prevalence of three important *Pantoea* species in these eleven African countries.

## Discussion

Bacterial infections by *Pantoea spp*. are recognized as being responsible for several diseases of plants, including important crop plants such as rice, maize, sorghum, onion and melon [34–43]. BB of rice caused by species of *Pantoea* were reported in several countries and include Benin, Togo, Korea, India, Australia, China, Italy, Venezuela, and Russia [10,11,40,44–49].

Given the fact that more than 25 species of *Pantoea* are currently known and among them several species can infect plants, efficient diagnostic tools are highly demanded by plant pathologists and extension workers. Some plant diseases were attributed to only three species of *Pantoea*, namely *P. agglomerans*, *P. ananatis* and *P. stewartii*, which can therefore be considered as the major *Pantoea* species infecting plants. For their diagnosis, several PCR methods are available and have been used but some of them produced amplicons with others species as well [14,50,51], while others are not well reproducible or are inaccessible in typical sub-Saharan laboratory due to specific equipment requirements and/or high costs of some reagents [14,17,18]. Notably, most assays target only one *Pantoea* species or subspecies. For instance, being of major concern, *P. stewartii* subsp. *stewartii* causing Stewart’s bacterial wilt can be detected by several methods but none of them can at the same time identify other bacteria of the genus *Pantoea* [14,16,18,52,53]. To the best of our knowledge, no robust diagnostic scheme exists that can specifically detect all three major *Pantoea* species that infect plants.

Based on whole genome sequences, we developed a new multiplex PCR scheme that can specifically detect the three major species of plant-pathogenic *Pantoea*, *P. agglomerans*, *P. ananatis* and *P. stewartii*. Different strategies can be followed when developing such a multiplex scheme using available whole-genome sequences. One possibility is to automatize the procedure by identifying genomic regions that are shared among a set of strains (e.g. the target species) and which are absent in another set of strains (non-target species). For instance, such an approach was used for the development of a *Xanthomonas oryzae*-specific multiplex PCR scheme that can differentiate the two pathovars *oryzae* and *oryzicola* [9]. The problem with this approach is that it might identify non-essential, often hypothetical genes as targets for the primer design. While present in the training set, it is hard to predict if these non-essential genes will be present and conserved in other, hitherto uncharacterized strains, especially when they originate from other geographical zones and/or belong to more distant genetic lineages.

Here, we targeted housekeeping genes, which are conserved throughout the genus, and relied on lineage (species)-specific sequence polymorphisms. This approach is considered as very robust but it cannot be ruled out that recombination events among strains from different species could undermine the universality of these primer pairs. Yet, we did not find any evidence for such events in any of the sequenced *Pantoea* strains that were analysed, including environmental isolates and strains isolated from human and plant samples. Nevertheless, because this study was focused on isolates from African rice leaves and seeds and only included a few reference strains from other continents (Additional file 1), it might be of interest to evaluate the new multiplex PCR tool on *Pantoea* strains isolated from other organisms (others plants, insects, other animals, humans) and from environmental samples.

To reduce the costs and handling time, we generated a multiplex PCR scheme that can work with both purified genomic DNA or with bacterial lysates. In both cases, sufficient specificity and sensitivity were obtained allowing detection of as low as 0.5 ng of DNA or 10^4^ CFU/mL for all three *Pantoea* species. Such a simple scheme will be of specific interest to phytopathologists, especially in Africa and other less-developed regions. Indeed, diseases due to infections by *Pantoea* appear to emerge in Africa as recently documented for Benin and Togo [10,11]. In this study, the presence of the three major plant-pathogenic *Pantoea* species has been demonstrated for eleven African countries. The fact that most of the BB-like symptomatic rice samples proved to contain a high number of *Pantoea* bacteria suggests that infection by *Pantoea* is an underestimated source for BB symptoms and might be widespread in Africa. However, more rigorous sampling schemes are required to determine the incidence and prevalence of *Pantoea* in various rice-growing areas in Africa.

Among the 609 *Pantoea* isolates, we detected 220 strains (36.1%; additional file 1) of *Pantoea* sp. that could not be assigned to any of the three species that are specifically targeted by the multiplex PCR scheme. This is an interesting observation that shows that the genus-specific primer pair does not only serve as an internal positive control of the multiplex scheme but that it has its own diagnostic value. Obviously, other species of *Pantoea* are present in Africa and are likely to cause disease of rice plants as well. Yet, it is still unknown whether or not this group of isolates contains other rice pathogenic species. Pathogenicity assays need to confirm or disprove their status as novel pathogens. Future work will address these isolates, using MLSA and whole genome sequencing.

While screening a large collection of bacterial isolates from rice samples, we also found strains that neither belonged to *Pantoea* nor to *Xanthomonas* (data not shown). Some of them were *Sphingomonas* strains [12], while others may represent new species and genera, which have so far not been connected to rice diseases. These isolates will be further studied by 16S rRNA analysis. From this study, it was concluded that the number of bacterial species that affect rice plants in Africa is certainly larger than previously thought.

## Conclusion

A new multiplex PCR scheme was developed to diagnose plant-pathogenic *Pantoea* spp. This tool enabled the efficient confirmation of the presence of *Pantoea* species (*P. ananatis* and *P. stewartii*) in Benin and Togo, as reported previously, and in several other African countries (Burkina Faso, Burundi, Ghana, Ivory Coast, Mali, Niger, Nigeria, Senegal, Tanzania). Moreover, we found evidence for the presence of *P. agglomerans* and other species of *Pantoea* on rice samples from several African countries. This new diagnostic tool will be very useful for crop protection services.

## Declarations

### Ethics approval and consent to participate

Not applicable.

### Consent for publication

Not applicable.

### Availability of data and material

Not applicable.

### Authors’ contributions

KK and RK conceived and designed the experiments. KK, SD, RA, RD evaluated the primers and multiplex PCR scheme by screening African strains. KK, RK and DS wrote the manuscript. All authors read and approved the final manuscript.

## Acknowledgements

We thank Toyin Afolabi (Africa Rice Center Cotonou, Benin), Sandrine Fabre and Florence Auguy (IRD, Cirad, University Montpellier, IPME, Montpellier, France) for excellent technical support. We are grateful to Charlotte Tollenaere for testing this new diagnostic tool in the international laboratory “LMI Patho-Bios” (IRD-INERA Observatoire des Agents Phytopathogènes en Afrique de l’Ouest) in Burkina Faso and for helpful comments on the manuscript.

## Authors’ information

KK, Institut de Recherche pour le Développement, Montpellier, France; Université de Montpellier, France & Africa Rice Center, Plant Pathology, Cotonou, Benin;

RA, Africa Rice Center, Plant Pathology, Cotonou, Benin;

RD, Africa Rice Center, Plant Pathology, Cotonou, Benin;

DS, Africa Rice Center, Plant Pathology, Cotonou, Benin;

RK, Institut de Recherche pour le Développement, Montpellier, France.

## Additional files

**Additional file 1**: List of bacterial strains used to evaluate the multiplex PCR scheme.

